# Learning a Pairwise Epigenomic and Transcription Factor Binding Association Score Across the Human Genome

**DOI:** 10.1101/2024.12.19.629547

**Authors:** Soo Bin Kwon, Jason Ernst

## Abstract

Identifying pairwise associations between genomic loci is an important challenge for which large and diverse collections of epigenomic and transcription factor (TF) binding data can potentially be informative. We therefore developed Learning Evidence of Pairwise Association from Epigenomic and TF binding data (LEPAE). LEPAE uses neural networks to quantify evidence of association for pairs of genomic windows from large-scale epigenomic and TF binding data along with distance information. We applied LEPAE using thousands of human datasets. We present evidence using additional data that LEPAE captures biologically meaningful pairwise relationships between genomic loci and expect LEPAE scores to be a resource.

## Background

Studying genomic loci in the context of pairwise relationships can be informative about the individual loci as well as the larger genomic regions in which they lie^1–4^. One prominent example of pairwise relationships used to study genomic loci is chromatin interactions identified by assays such as Hi-C^5–7^. While informative, such assays miss relationships associated with physical interactions beyond the detection thresholds of the assays as well as relationships that are not linked with physical interactions. For example, two loci might be associated in that they are part of the transcribed region of the same gene but do not physically interact.

An alternative strategy to identify pairwise relationships between loci has been based on correlating epigenomic assay signals at distal loci directly or with gene expression across cell types, often focusing on linking enhancers with target genes^2,4,8,9^. While correlations based on epigenomic mark signals have also been informative, they are limited in the use of available information in large and diverse compendia of genome-wide maps of histone modifications and variants, chromatin accessibility, chromatin state annotations, and transcription factor (TF) binding and thus are likely limited in the types of associations they capture. Correlation-based measures also cannot effectively reveal associations between two distal loci that have constitutive activity.

To address these issues, we developed a method, Learning Evidence of Pairwise Association from Epigenomic and TF binding data (LEPAE), which systematically identifies pairwise associations between pairs of genomic windows by leveraging a large compendium of epigenomic and TF binding data from diverse assays and cell types along with genomic distance information. LEPAE assumes that pairs of windows at specific distances apart are more likely to have associations reflected in epigenomic and TF binding data than random pairs of windows. Based on this assumption, LEPAE trains a neural network classifier with a Siamese architecture to distinguish pairs of windows at fixed distances apart from randomly mismatched pairs of the same set of windows. LEPAE then applies the trained classifier to score evidence of association for pairs of windows across a genome. LEPAE’s approach is an adaption of the ‘Learning Evidence of Conservation from Integrated Functional genomics data’ (LECIF) method, which we previously used to score evidence of conservation at the functional genomics level between human and mouse^10^. Additionally, LECIF or extensions of the framework have been applied for comparisons between human and pig^11^ and in the context of plant genomics^12^. We apply LEPAE to score pairs of windows in the human genome up to 1 Mb apart. Using gene annotations, topological associated domains, chromatin state annotations, and genome-wide association study (GWAS) variants, we validate that the LEPAE score reflects associated or similar properties in pairs of loci. We expect the LEPAE score to be a complementary resource to existing genomic annotations and datasets for understanding relationships among genomic loci.

## Results

### Overview of LEPAE

LEPAE quantifies evidence of association between pairs of binned genomic windows based on epigenomic and TF binding data. LEPAE does this for each pair of windows at a resolution of *r* bases within *D* bases. To do this, LEPAE first trains classifiers for each pairwise distance *d*, where *d* ranges from *r* to *D* bases at increments of *r* bases. The classifiers are trained to discriminate pairs that are *d* bases apart as positive pairs from a negative set of pairs. The negative pairs are randomly mismatched positive pairs within each chromosome, which allows for controlling for chromosome-specific characteristics. LEPAE assumes that *D* is substantially less than the chromosome length and positive pairs relative to negative pairs more likely represent associated windows given their proximity.

LEPAE trains each classifier using epigenomic and TF binding data as features for the classification. The specific type of classifier LEPAE uses is a Siamese neural network^13^ (**Fig. 1**; **Methods**). A similar classifier was previously shown to be effective in the LECIF method for learning patterns from two input feature vectors^10^. The Siamese neural network architecture forces a classifier to learn the same weights for both windows in each input pair and thus its predictions are invariant to how the two windows are ordered when provided as input to the classifier.

**Figure 1.**
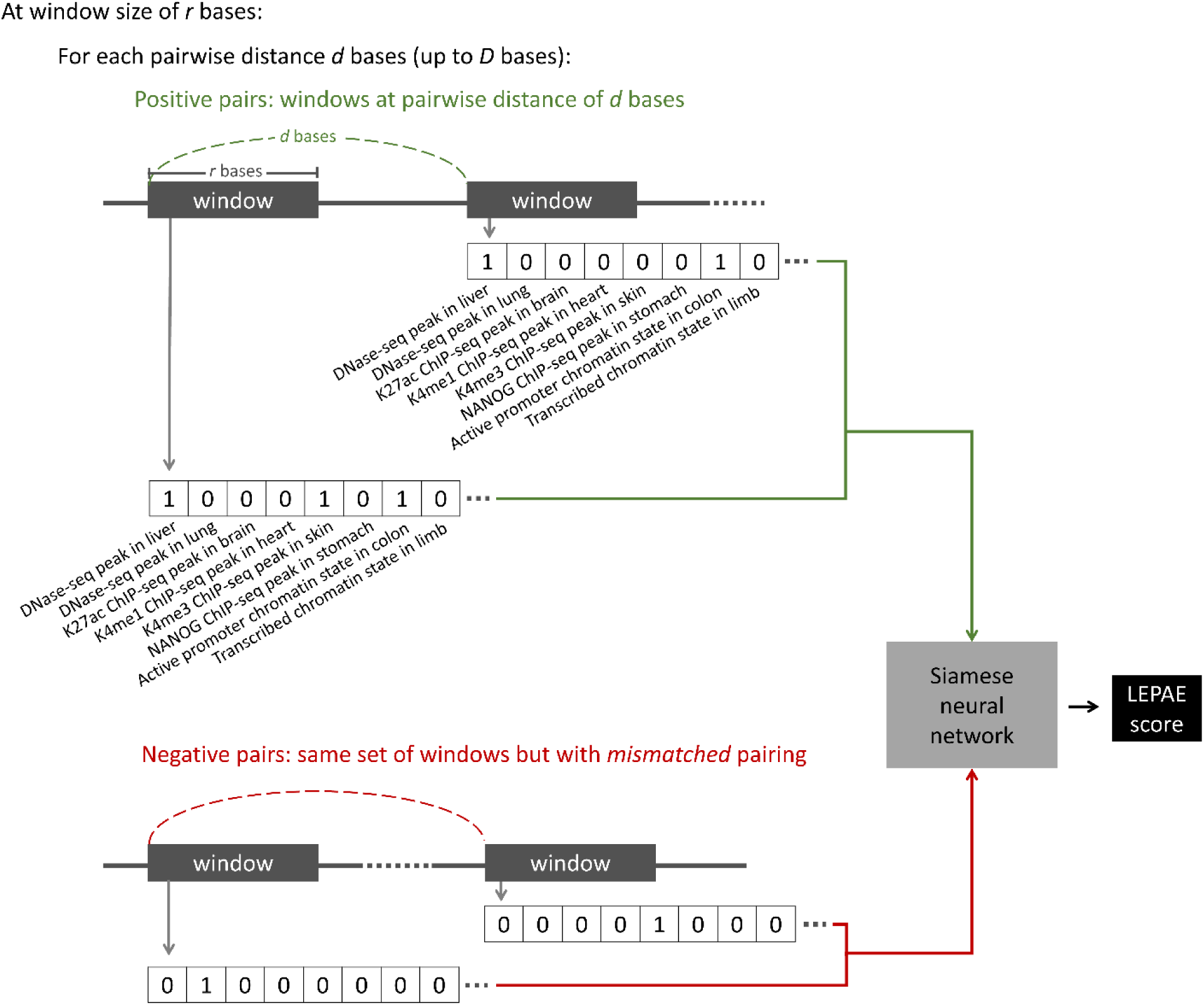
Overview of the LEPAE method. For every genomic window of *r* bases, LEPAE generates a feature vector from thousands of human epigenomic and TF binding annotations. Only a subset of the features is shown here. A pair of two such feature vectors are provided as input to a Siamese neural network. At each pairwise distance of *d* bases, the classifier is trained to distinguish positive pairs (green), which are genomic windows that are *d* bases apart, from negative pairs (red), which are randomly mismatched pairs of the same set of windows. To generate the LEPAE score for a pair of windows that are *d* bases apart and held out from training, LEPAE ensembles predictions from multiple classifiers trained on random subsets of training data and also for pairwise distances of *d – r, d,* and *d + r* (**Methods**). Feature labels (e.g. DNase-seq peak in liver) are not provided to LEPAE.

For each pair, LEPAE uses two binary feature vectors corresponding to epigenomic and TF binding data overlapping the pair’s two genomic windows. In our application, the binary features correspond to whether a window overlaps with peak calls from DNase-seq experiments or ChIP-seq experiments of histone modifications, histone variants, or TFs or individual ChromHMM chromatin state annotations^14,15^. These peak calls and chromatin state annotations cover a diverse set of human cell and tissue types generated by the ENCODE^16^ or Roadmap Epigenomics Projects^17^ (**Fig. 1**). Because the same set of windows are used in positive and negative pairs, each classifier learns pairwise characteristics of epigenomic and TF binding data from the two windows in a pair as opposed to characteristics of marginal features. During training, chromosomes for which LEPAE will make predictions are held out (**Methods**). Once trained, each classifier makes predictions for pairs of genomic windows at its corresponding distance for chromosomes that were held out from training. LEPAE then ensembles individual classifier predictions to improve performance and consistency across different pairwise distances (**Methods**). In our application, we applied LEPAE for pairs of 1-kb human genomic windows with distances *d* ranging from 1 kb to 100 kb in increments of 1 kb (score resolution *r*=1 kb; maximum pairwise distance *D*=100 kb). In addition, we applied LEPAE for pairs of 10-kb genomic windows with pairwise distances *d* ranging from 10 kb to 1 Mb in increments of 10 kb (*r*=10 kb; *D*=1 Mb). As a result, we annotated more than 302 million and 30 million pairs of 1-kb and 10-kb genomic windows, respectively, with LEPAE scores (**Fig. 2, 3c; Supplementary Fig. 1c**). We illustrate LEPAE scores at two loci (**Fig. 2**), which highlights how LEPAE scores can be higher for pairs of loci within the same annotated genes despite making its predictions independent of such annotations.

**Figure 2.**
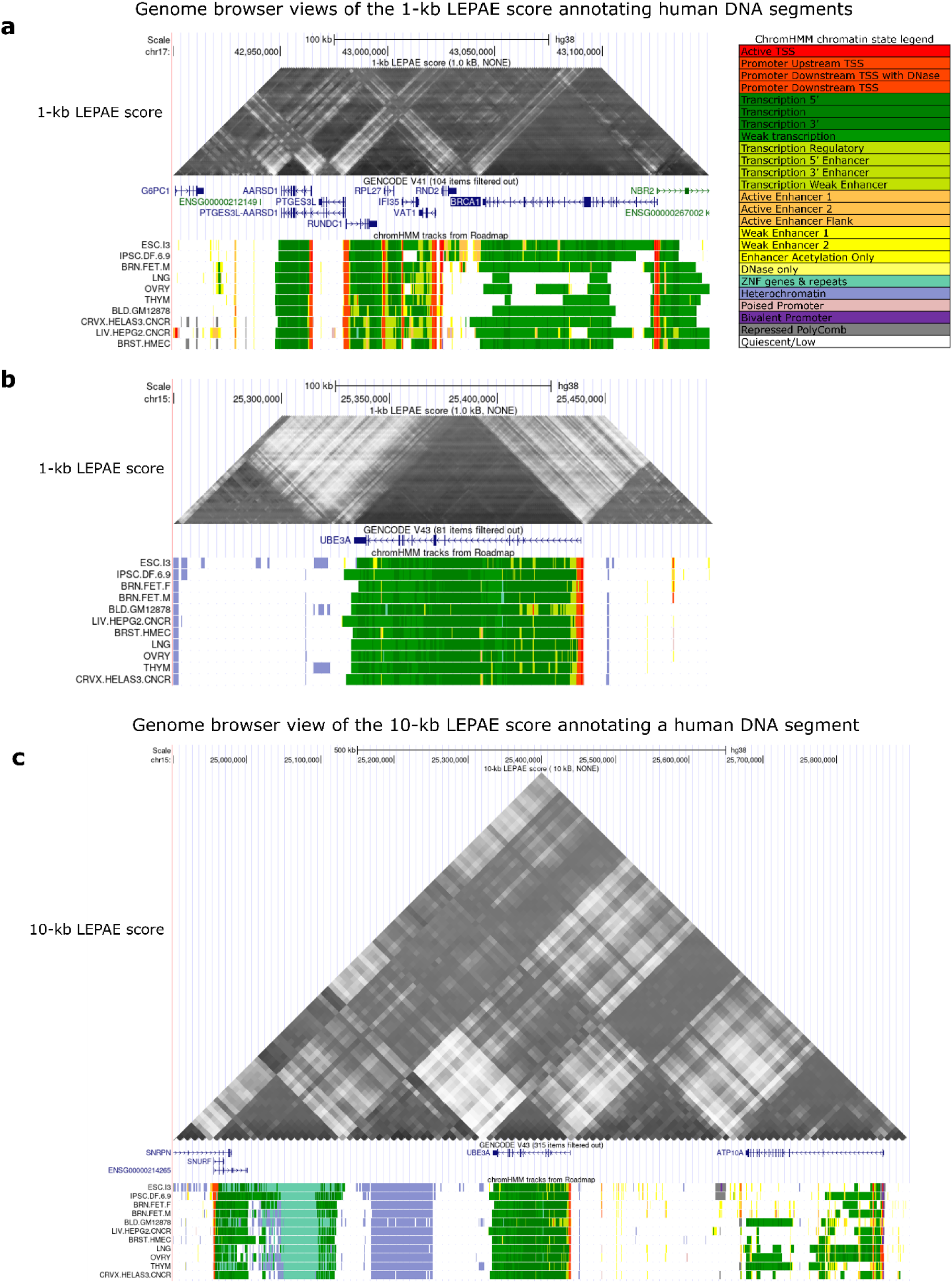
Genome Browser Views. **a.** UCSC Genome Browser^29^ view of the 1-kb LEPAE score annotating gene BRCA1 and its neighboring regions. The score is shown in the top as a heatmap in a format primarily used to visualize Hi-C contact matrices^30^. Darker colors correspond to higher scores. Below the score are GENCODE V41 gene annotations^23^, followed by ChromHMM chromatin state annotations^14,15^ for different epigenomes from the Roadmap Epigenomics Project^17^, which were provided as input. While chromatin state annotations from 127 epigenomes were used as input features, only a subset is shown here. State legend is on the top right. **b.** Same as **a** but for gene UBE3A instead of BRCA1 and not reshowing the chromatin state legend. **c.** Same as **b** showing gene UBE3A but for 10-kb LEPAE score instead of 1-kb.

### LEPAE’s predictive power and relationship to distance

LEPAE has strong predictive power when differentiating positive pairs from negative pairs, all held out from training, especially for pairs with small pairwise distances. For the 1-kb model, we observe the strongest predictive power for pairs with the smallest pairwise distance, 1 kb, with a mean area under the receiver operating characteristic curve (AUROC) of 0.92 (**Fig. 3a**). For pairs with larger distances, the LEPAE score’s predictive power declines, with a minimum mean AUROC of 0.72 at a pairwise distance of 99 kb. Similarly, we observe the greatest predictive power for the 10-kb model with an AUROC of 0.92 at its smallest distance, 10 kb, and with the predictive power declining to the lowest mean AUROC of 0.68 at its largest distance of 1 Mb (**Supplementary Fig. 1a**). Moreover, LEPAE achieves better predictive power than a score learned by decision trees instead of Siamese neural networks at all distances (**Fig. 3a; Supplementary Fig. 1a; Methods**).

**Figure 3.**
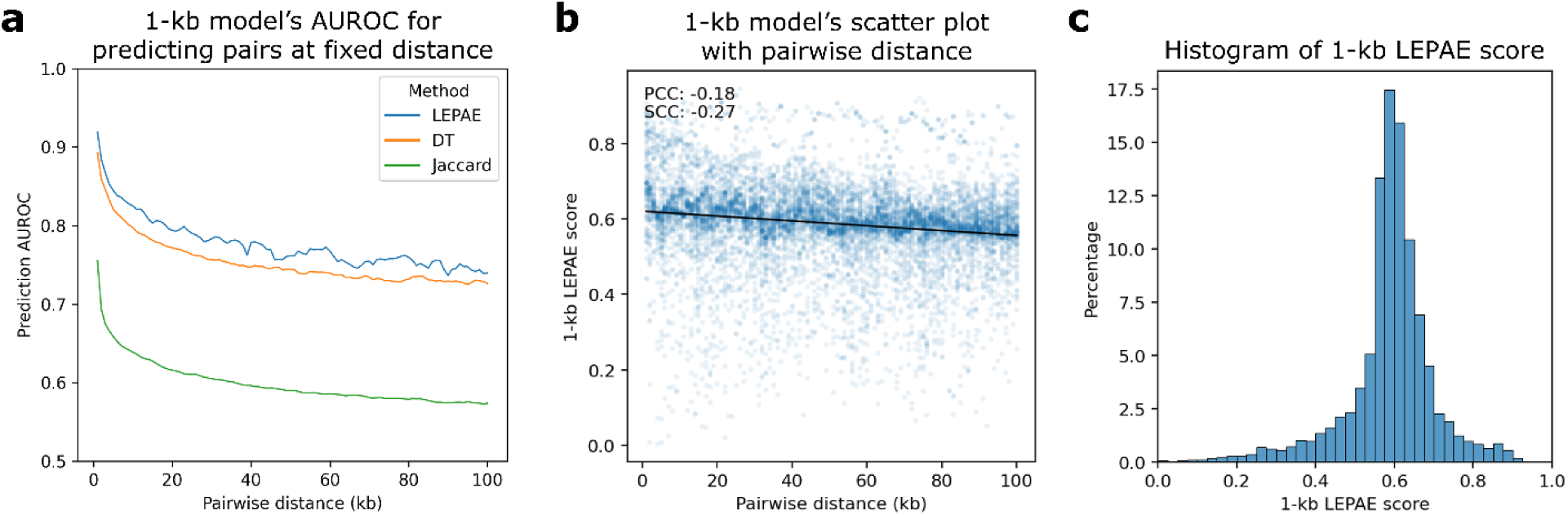
Characteristics of the 1-kb LEPAE score. **a.** Relationship between pairwise distance and prediction AUROC for 1-kb model. For each pairwise distance (x-axis), mean prediction AUROC of the 1-kb LEPAE score for distinguishing pairs of windows at the distance from randomly mismatched pairs of the same windows is shown in blue. The mean is computed from two sets of classifiers trained on non-overlapping training sets (**Methods**). Mean AUROC values are shown when a decision tree (DT) instead of a neural network was used as the supervised classifier are shown in orange (**Methods**). Mean AUROC values computed using Jaccard index instead of the LEPAE score to perform the same classification task are shown in green. Values belonging to the same method are connected by piecewise linear interpolation. **b.** Scatter plot showing with a blue dot for each pair of windows the pairwise distance (x-axis) and the 1-kb LEPAE score (y-axis). Ten thousand random pairs are shown. A linear regression line fitted to the ten thousand random pairs is shown in black. PCC and SCC, computed from 1 million randomly selected pairs, are shown in the top left. **c.** Distribution of the 1-kb LEPAE score. Forty bins ranging from 0 to 1 with increments of 0.025 were used. A 10-kb score resolution version of this figure is in **Supplementary Figure 1**.

We also confirmed that the LEPAE score is largely distinct from 1D genomic distance (**Fig. 3b; Supplementary Fig. 1b**). We observed only modest negative Pearson correlation coefficient (PCC) values of -0.18 and -0.26 for their correlations, when using the 1-kb and 10-kb LEPAE scores, respectively. Even at the maximum pairwise distances considered (*D*), 3% and 1% of pairs score above the 95th percentile among all scores at 1-kb and 10-kb resolution, respectively. These results indicate that LEPAE can highlight pairs of windows that are distal but exhibit sufficient evidence of association based on their epigenomic and TF binding data.

To confirm that LEPAE learns information beyond similarity in input features, we compared the LEPAE score to Jaccard index, which was computed for every pair of windows to which LEPAE was applied (**Methods**). Specifically, for each pairwise distance we compute the correlation between the LEPAE score and Jaccard index for pairs of windows with that pairwise distance. Consistent with LEPAE learning information beyond similarity in input features, we observed only a moderate correlation with the Jaccard index, with a mean PCC of 0.16 and 0.39 at 1-kb and 10-kb resolutions, respectively (**Supplementary Fig. 2**). For the task of differentiating positive pairs from negative pairs, LEPAE achieves consistently better predictive performance than the Jaccard index at both resolutions (**Fig. 3a; Supplementary Fig. 1a**). These results confirm that LEPAE captures information beyond similarity in features.

### High-scoring pairs highlight associated loci based on annotation of genes and chromatin states

We next analyzed the LEPAE score in the context of gene annotations, which were not used in learning the score. Specifically, we asked whether the LEPAE score was higher for pairs of bases that fell within the gene body of the same gene compared to a matched control set. The matched control set consisted of pairs of bases for the same set of genes and the same distance apart but with one base inside and the other outside the gene body. For these control pairs, the region between the two bases crosses the gene boundary (**Methods**). We found that pairs of bases within the gene body have higher 1-kb resolution LEPAE score than pairs that crossed gene boundaries (**Fig. 4**; Wilcoxon signed-rank test *p*<0.0001; **Methods**). In addition, we found that this difference is stronger than the difference observed with Hi-C data and Jaccard index (**Fig. 4**; Mann-Whitney U test *p*<0.0001; **Methods**).

**Figure 4.**
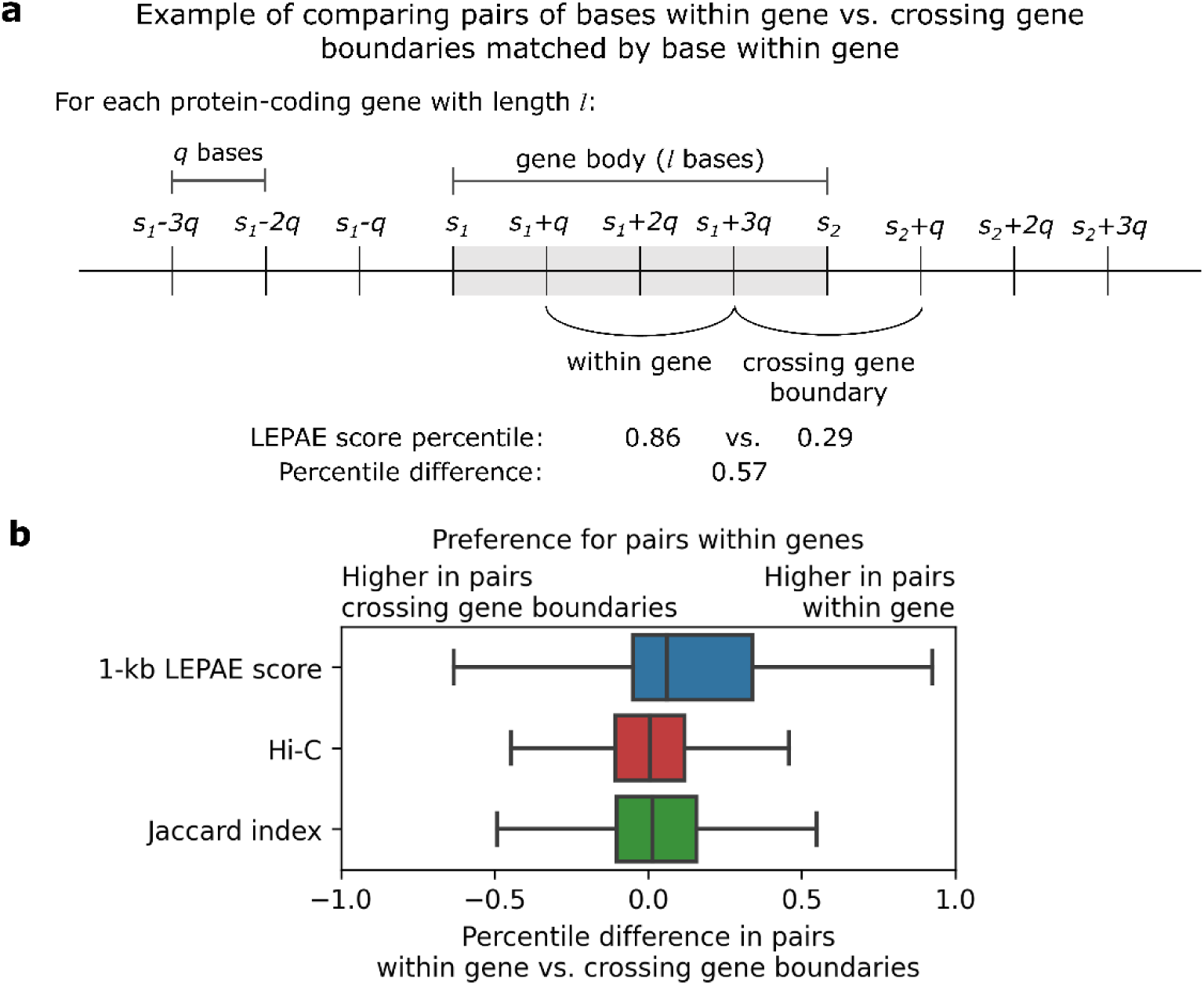
1-kb LEPAE score’s relationship to genic annotation. **a.** For a protein-coding gene of length *l*, its quartile length *q* is used to define bases within the gene body, spanning from the transcription start site (TSS) *s_1_* to transcription end site (TES) *s_2_*, and upstream and downstream of the gene body. A pair of bases within a gene (e.g*. s_1_+q* and *s_1_+3q*) is compared to a pair of bases crossing the gene boundary (e.g. *s_1_+3q* and *s_2_+q*), where the two pairs are matched by their pairwise distance (e.g. *2q*) and share one base located within the gene (e.g. *s_1_+3q*). The percentile difference is computed by taking the LEPAE score percentile of the former and subtracting it by the score percentile of the latter. This approach was repeated for normalized Hi-C contact frequency and Jaccard index for comparison (**Methods**). **b.** Shown for the 1-kb LEPAE score, normalized Hi-C contact frequency, or Jaccard index is the distribution of percentile differences between pairs with both bases within a gene and pairs with only one base within a gene. A positive percentile difference indicates a preference for pairs within genes over pairs crossing gene boundaries.

To better understand the nature of pairs of windows that received a high LEPAE score, we also analyzed the score for pairs of genomic elements using a universal chromatin state annotation^18^ (**Methods**). This annotation provides a single chromatin state assignment per genomic position based on over 1000 human epigenomic datasets from more than 100 cell or tissue types. In total there were 100 states in this annotation that were manually grouped into 16 broader groups. This chromatin state annotation was not directly provided to LEPAE as input, though we note that there is overlap in the input datasets provided to LEPAE and the datasets used to learn the chromatin state model. We first computed the percentile of 1-kb LEPAE score for all pairs of windows with pairwise distances of 5 kb and 50 kb, separately, and then the mean percentile for each pair of chromatin states for both pairwise distances (**Methods, Fig. 5; Supplementary Fig. 3,6a**). For the 5kb distance, pairs of the same state had a high mean 1-kb LEPAE score percentile with an average of 0.69, compared to 0.51 for any two states with that pairwise distance (**Fig. 5a,d; Supplementary Fig. 3,6a**). Notably, pairs of transcription-associated states (Tx, TxEnh, TxEx) with a pairwise distance of 5 kb had notably high mean LEPAE score percentiles ranging from 0.84 to 0.96 (**Fig. 5a**). Similarly, pairs of active promoter-associated states (PromF, TSS) had a mean 1-kb LEPAE score percentile ranging from 0.63 to 0.87 (**Fig. 5a**). We saw consistent trends using a pairwise distance of 50 kb with pairs of the same state scoring 0.66 on average compared to an overall mean of 0.54 for all state pairs that are 50 kb apart.

**Figure 5.**
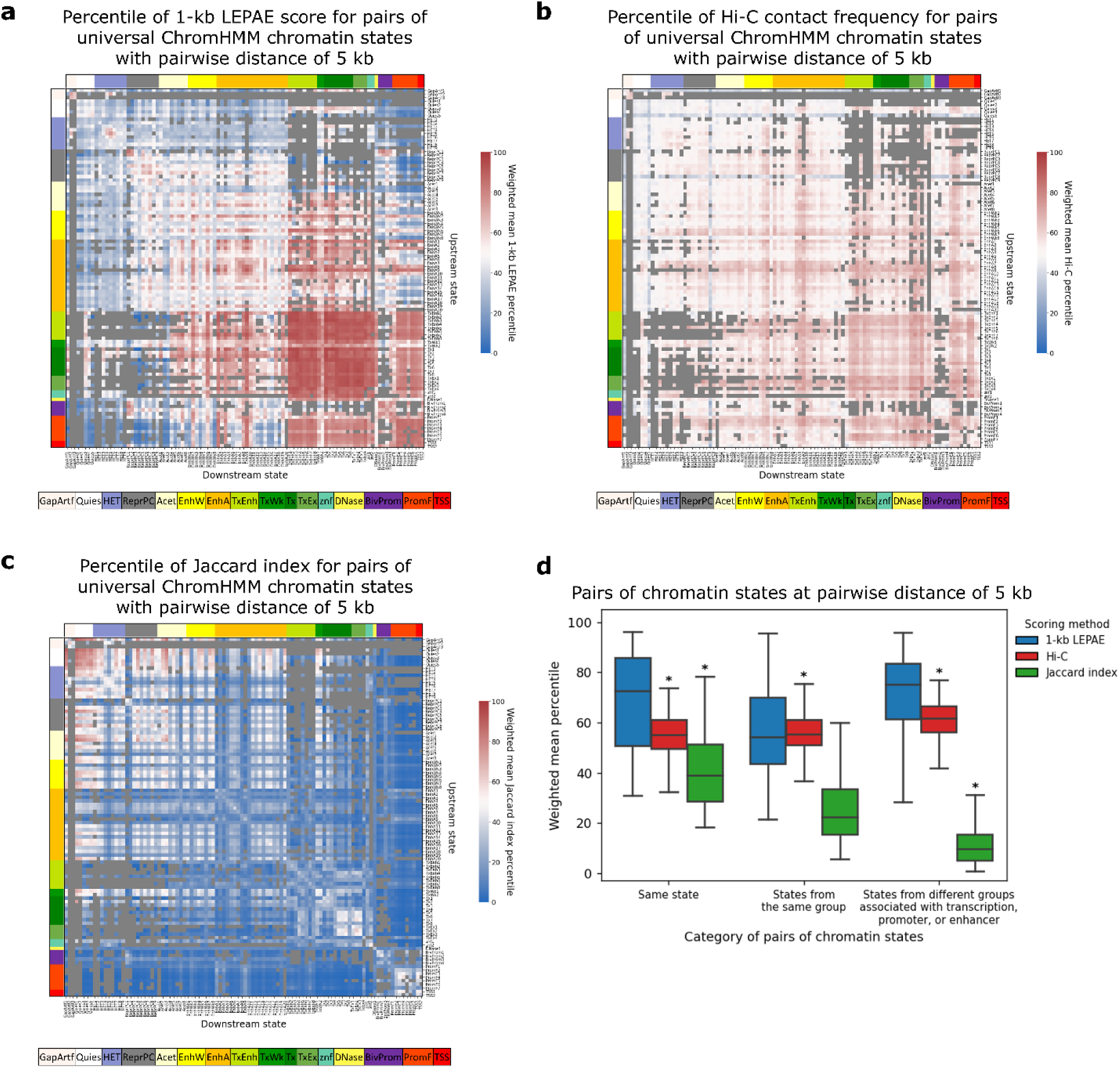
1-kb LEPAE score’s relationship to pairs of chromatin states with pairwise distance of 5 kb. **a.** Each cell in the heatmap corresponds to a state pair, one annotating the upstream window of a pair of 1-kb windows (row) and the other annotating the downstream 1-kb window of the same pair (column) with their pairwise distance fixed to 5 kb. The states are from a universal chromatin state annotation based on more than 1000 epigenomic datasets from more than 100 cell or tissue types^18^. The ordering of states in the rows and columns are the same. Color shown next to the topmost row or leftmost column corresponds to the state group of each state along the column or row, respectively, according to the legend on the bottom. The colors shown in the cells correspond to a weighted mean LEPAE score percentile of pairs of windows that are 5 kb apart and are annotated by the states specified in the row and column (**Methods**). Color legend for the score is shown on the right. Pairs of states that together cover less than 0.0005% of all the bases annotated by pairs of states were excluded from the analysis and grey is shown in their corresponding cells. **b.** Similar to **a** but with the color in each cell corresponding to a weighted mean Hi-C contact frequency percentile. **c.** Similar to **a** but with the color in each cell corresponding to a weighted mean Jaccard index percentile. **d.** Shown for three different categories of pairs of ChromHMM chromatin states are the distribution of the weighted mean percentile of the 1-kb LEPAE score, Hi-C contact frequency, or Jaccard index, at a fixed pairwise distance of 5 kb as done in **a-c**. Among the three categories, the first category shown on the leftmost position on the x-axis corresponds to pairs of the same state. The second, shown in the middle, corresponds to pairs of different states belonging to the same state group. The last category shown on the rightmost position corresponds to pairs of states where both states are associated with active promoters, transcription, or enhancers but are from different state groups. These three categories of pairs of states do not overlap with each other. Within each category, the distributions of the weighted mean percentile of the 1-kb LEPAE score, Hi-C contact frequency, and Jaccard index are shown in blue, red, and green, respectively, according to the score legend on the bottom. An asterisk above a distribution denotes that there is a significant difference between it and the distribution of weighted 1-kb LEPAE score percentiles for the same category based on a Mann-Whitney U test (p-value < 0.0001). Similar versions of the figure panels but for pairwise distance of 50 kb and also for 10-kb score resolution at pairwise distances of 50 kb and 500 kb are in **Supplementary Figures 3-6**.

We also observed cases of pairs of different types of states having a high mean 1-kb LEPAE score. For instance, at a pairwise distance of 5 kb, pairs of states which are associated with active promoters, transcription, or enhancers, but did not belong to the same state group scored high with a mean LEPAE score percentile of 0.70 on average, close to the average of 0.69 for pairs of the same state. The state groups were previously defined in Ref. ^18^ based on model parameters and state enrichments for external genome annotations. Notably, a state associated with enhancer activity in blood and thymus (EnhA9) scored high when paired up with transcription-associated states, such as a state associated with transcription and enhancer activity in blood (TxEnh2) or an active transcription state (Tx1), with the mean 1-kb LEPAE score percentile of 0.92 for both at a pairwise distance of 5 kb (**Fig. 5a**). This was similar to the mean 1-kb LEPAE score percentile of 0.90 when enhancer state EnhA9 state was paired with itself at that pairwise distance. This state also scored high when paired with active promoter chromatin states (PromF1-7, TSS1-2) with the mean LEPAE score percentile ranging from 0.79 to 0.87. Also at distance 50 kb, pairs of states that were previously characterized as being active promoter, transcription, or enhancer associated and belong to different state groups scored high with a mean 1-kb LEPAE score percentile of 0.72 on average, close to the average of 0.66 for pairs of the same state (**Supplementary Fig. 6a**). These results suggest that the LEPAE score captures biologically meaningful relationships between loci that can be associated with different annotations.

We further observed pairs of states that had markedly low mean LEPAE scores. For example, we saw that the quiescent states (Quies1-5) tended to score low when paired up with states associated with transcription or enhancer activity. As a specific example, at a pairwise distance of 5 kb, pairs of a quiescent state (Quies5) and a state characterized by transcription and enhancer (TxEnh2) had a mean 1-kb resolution LEPAE score percentile of 0.04 (**Fig. 5a**). This suggests that the LEPAE score can also highlight pairs of chromatin states that are less likely to be associated.

We also investigated whether these patterns captured by the 1-kb LEPAE score are recapitulated with Hi-C contact frequency or Jaccard index (**Fig. 5d**, **Supplementary Fig. 3,6a**; **Methods**). We observed that the 1-kb LEPAE score better highlighted pairs of the same state, states from the same state group, and states associated with active promoters, transcription, or enhancers that belong to different state groups (Mann-Whitey U test *p*<0.001). For those same set of state pairs, Hi-C contact frequency tended to be close to the median, not differentiating any particular group of state pairs from the rest. On the other hand, Jaccard index at 1-kb resolution scored pairs of the same state high and scored pairs of states that are associated with active promoters, transcription, or enhancers and belong to different groups low, which is expected from the similarity-based nature of the index. We did not observe the patterns captured by the 1-kb resolution scores with 10-kb resolution scores (**Supplementary Fig. 4-6**). However, we were not necessarily expecting to since the chromatin states are defined at 200-bp resolution, while the much larger 10-kb windows tend to include more states, making it harder to attribute a high score observed in a pair of windows to a specific pair of chromatin states. Overall, these results suggest the LEPAE score can highlight not only similar loci but also loci exhibiting distinct but associated properties.

### Relationship to chromatin contact frequency

We next studied the relationship between LEPAE scores and chromatin interaction data. Specifically, we compared the 1-kb LEPAE score to normalized 1-kb resolution chromatin interaction frequencies collected in a Hi-C experiment in GM12878 (**Methods**). We computed the PCC between the LEPAE score and a normalized Hi-C matrix for pairs with the same pairwise distance and observed a low mean PCC (0.06). We also observed similar results when comparing 10-kb LEPAE score to 10-kb resolution Hi-C data (mean PCC of 0.06). These results suggest that the LEPAE score captures associations largely distinct from Hi-C chromatin interaction frequencies.

We also related the LEPAE score to an annotation of topologically associating domains (TAD) from the same Hi-C experiment (**Methods**). Among pairs of windows at a particular pairwise distance, we compared pairs within the same TAD to pairs within the same distance that cross a TAD boundary, with one window inside a TAD and the other outside the TAD. For the 1-kb LEPAE score, at 93 out of 100 pairwise distances, pairs within TADs had a significantly higher score distribution than those crossing TAD boundaries (**Fig. 6**; Mann-Whitney U test *p*<0.0001; **Methods**). We observed similar results with the 10-kb LEPAE score, with statistically higher score distribution for pairs within TADs at all pairwise distances (**Supplementary Fig. 7**). While we observed a low overall correlation between LEPAE scores and Hi-C contact frequencies, LEPAE scores do show some level of agreement with Hi-C data as reflected in the TAD analysis.

**Figure 6.**
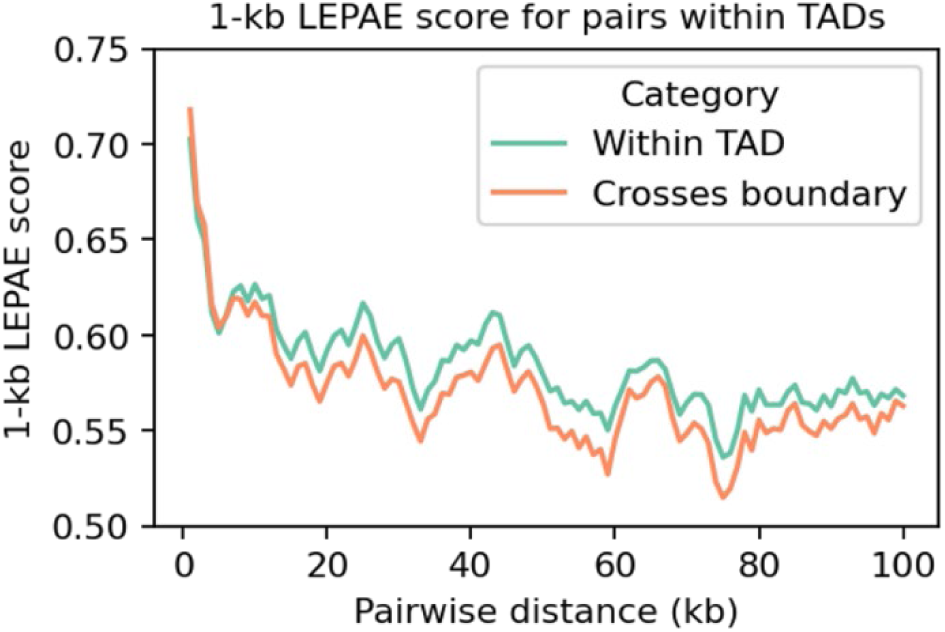
1-kb LEPAE score’s relationship to Hi-C contact frequency. Shown for each pairwise distance (x-axis) is the mean 1-kb LEPAE score for pairs of windows located within a topologically associating domain (TAD) (turquoise) or the mean score for pairs of windows crossing a TAD boundary (peach). Values belonging to the same category are connected by piecewise linear interpolation. A 10-kb resolution score version of this figure is in **Supplementary Figure 7**.

### Relationship to fine-mapped GWAS variants

We next analyzed whether pairs of variants that were fine-mapped to be a causal variant for the same trait at the same loci had a higher LEPAE score than matched control pairs. We did this analysis for a set of 94 UK Biobank traits that were previously fine-mapped using the SuSiE^19^ and FINEMAP^20^ methods^21,22^. For each actual pair of fine-mapped variants, we extracted two matched control pairs of windows selected such that each control pair shared one window with the original pair and the two windows of the control pairs were the same distance apart as the original pair. As a result, all control pairs had a fine-mapped variant in one window. We conducted the analysis at 1-kb resolution for all pairs within 100 kb and at 10-kb resolution for all pairs within 1 Mb. Of the 94 traits, 73 to 78 traits had a higher LEPAE score for the original pairs compared to the control pairs with the specific number of traits varying with the fine-mapping method and score resolution (**Fig. 7**). In comparison, only 47 would be expected by chance (*p*<10^-7^). Also in comparison, 34 to 62 traits had a higher Jaccard index for the original pairs than control pairs, and 66 to 71 traits for Hi-C contact frequency (**Fig. 7**). These results suggest that pairs of loci with higher LEPAE scores may be more likely to share phenotypic associations.

**Figure 7.**
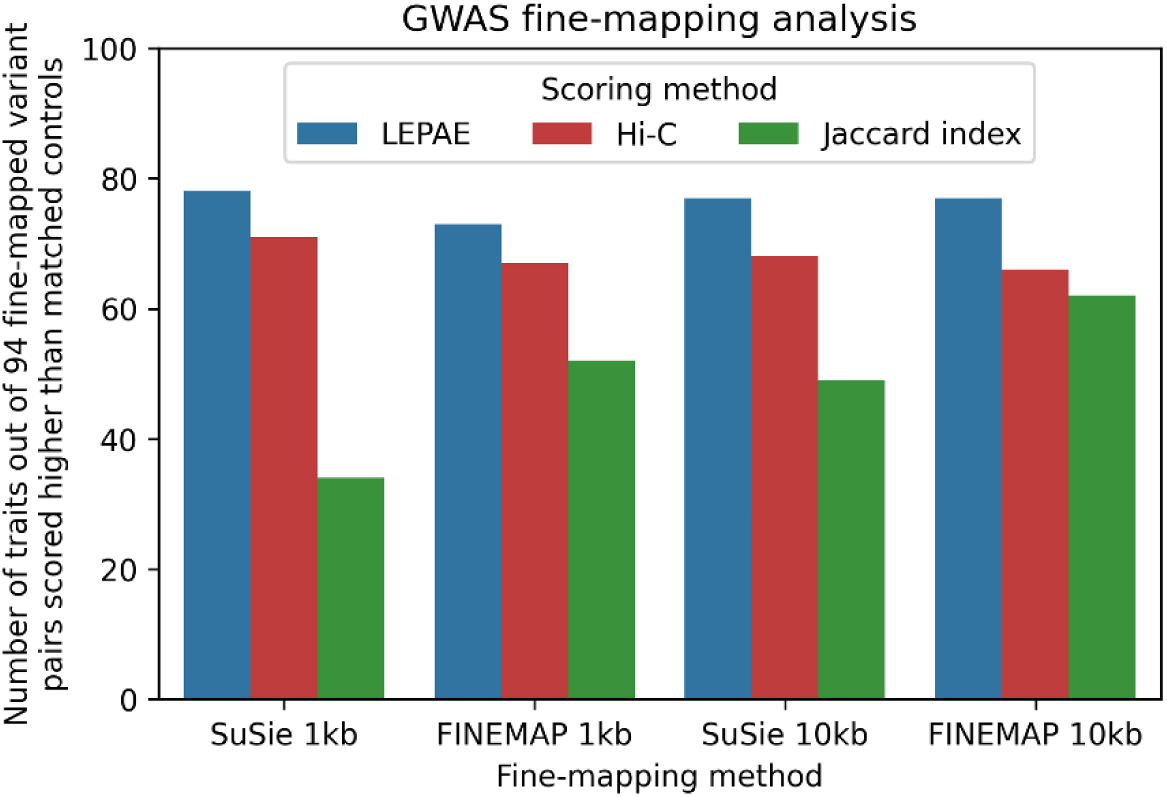
LEPAE’s relationship to fine-mapped GWAS variants. The figure shows out of 94 UK biobank traits^21,22^ for how many the LEPAE score was higher for pairs of fine-mapped variants than matched control variants. The results are shown for 1-kb resolution, which considers pairs up to 100-kb apart, and 10-kb resolution, which considers pairs up to 1 Mb apart, for both the SuSiE and FINEMAP methods. For comparison, results are also shown for normalized Hi-C contact frequency and the Jaccard index.

## Discussion

Here we presented LEPAE, a method that scores evidence for pairwise association between genomic windows based on a large collection of functional genomic datasets. LEPAE is a variant of LECIF^10^, a method to score evidence of conservation between two species based on functional genomics data. In contrast, LEPAE scores evidence of association based on functional genomics data for pairs of loci within the same species.

While the results we presented suggest LEPAE will be informative for studying pairwise relationships between genomic loci, it does have limitations. As the score characterizes relationships at fixed distances, it is not applicable to studying interchromosomal associations. Also, the score does not directly provide information about relationships involving more than two pairs of loci.

While here we applied LEPAE at 1-kb and 10-kb resolutions with a maximum distance of 1 Mb, we could in the future aim for a finer resolution and larger pairwise distances at a higher computational cost. When training LEPAE, here we assumed the maximum pairwise distance *D* was substantially less than the chromosome length and restricted negative pairs to consist of windows from the same chromosome to control for chromosome-specific characteristics during classification. An avenue for future investigation is to extend LEPAE for predictions at substantially larger distances by removing the restriction that the windows in negative pairs come from the same chromosome. Moreover, although here we applied LEPAE to epigenomic and TF binding data from the human genome, future work could investigate applying LEPAE with additional types of functional genomics data such as RNA-seq data and also to other widely studied species such as mouse or rat. We expect our existing LEPAE scores as well as the approach applied to other data will be a resource for studying relationships between genomic loci.

## Conclusions

We applied LEPAE with more than 3000 functional genomic datasets including maps of chromatin accessibility, histone modifications and variants, transcription factor binding, and chromatin state annotations from various tissue and cell types. The resulting LEPAE scores had greater predictive power for differentiating pairs of windows at a fixed distance from randomly mismatched pairs of the same windows than a baseline approach of computing the Jaccard index of input features. Using external annotations not provided as input features, including gene annotations, fine-mapped variants from GWAS, and TADs, we presented analyses suggesting LEPAE scores reflect biologically meaningful associations. We envision the LEPAE scores to serve as a resource for examining pairwise relationships across the genome.

## Data availability

The LEPAE scores are available in the LEPAE repository, https://github.com/ernstlab/LEPAE. Links to data files from the ENCODE^16^ or Roadmap Epigenomics Projects^17^ that were used to generate input features to LEPAE are listed in the same repository as well as in **Supplementary Table 1**. For analysis using gene boundaries, we used GENCODE annotation (V38)^23^. The in situ Hi-C data for GM12878^7^ used for the Hi-C analysis is experiment 4DNES3JX38V5^7^ on the 4DN Nucleome Data Portal^24^ and available from https://4dn-open-data-public.s3.amazonaws.com/fourfront-webprod/wfoutput/a98ca64a-861a-4a8c-92e9-586af457b1fb/4DNFI1UEG1HD.hic. TAD coordinates for the experiments are from the 3D Genome Browser^25^ and available from http://3dgenome.fsm.northwestern.edu/downloads/hg38.TADs.zip. The variants used in the fine-mapping analysis are available from https://www.finucanelab.org/data^21,22^. Universal chromatin state annotation was downloaded from https://github.com/ernstlab/full_stack_ChromHMM_annotations^18^.

## Code availability

The LEPAE software is available at the LEPAE repository, https://github.com/ernstlab/LEPAE.

## Methods

### Defining genomic windows

We binned all autosomal chromosomes and chrX into non-overlapping 1-kb windows and separately into non-overlapping 10-kb windows. We used hg38 as the genome assembly.

### Input features

Each pair of windows was assigned two feature vectors, one corresponding to the upstream window and the other corresponding to the downstream window. For each window, each peak call corresponded to a binary feature. If a genomic window overlapped a peak call in an experiment, the corresponding value in the feature vector was set to 1, otherwise it was set to 0. The chromatin state annotations were one-hot encoded such that each binary feature corresponded to the presence of a chromatin state in a cell or tissue type.

Peak calls for DNase-seq and ChIP-seq experiments of histone modifications, histone variants, and TFs were generated by ENCODE and processed by ENCODE4 pipelines^16^. ChromHMM chromatin state annotations^14^ were from the 25-state model learned from imputed data for 127 cell and tissue types from the Roadmap Epigenomics Project^15,17^ that were previously lifted over from hg19 to hg38 using the liftOver tool^26^.

### Pairwise distance

For 1-kb windows, we varied the pairwise distance from 1 kb to 100 kb with increments of 1 kb, resulting in 100 distances. For 10-kb windows, we varied the pairwise distance from 10 kb to 1 Mb with increments of 10 kb, resulting in another 100 distances. Distances were measured based on the first base of each window.

### Defining positive and negative pairs

For each pairwise distance *d*, pairs of windows with *d* bases between their first bases were defined as positive pairs. To generate negative pairs, we randomly shuffled the pairing of the positive pairs within the same chromosome, resulting in pairs of windows that are not necessarily *d* bases apart from each other but from another window in the same chromosome.

### Classifier

Each neural network had a Siamese architecture^13^ consisting of two identical sub-networks, which share their weights, followed by a final sub-network that combines the output from the two sub-networks to generate a final prediction. The two input feature vectors were fed into the two sub-networks. All sub-networks had a single hidden layer, resulting in two hidden layers in total.

To set the hyper-parameters of a classifier for a fixed pairwise distance *d*, window size *r*, and a particular set of training pairs, LEPAE conducted a random search, where it generated 10 classifiers, each with different randomly selected combination of hyper-parameters. Each classifier was trained on the same set of 50,000 training pairs and evaluated on the same set of 5,000 validation pairs. Hyper-parameters were varied as follows during random search:

- Batch size: 16, 32, 64
- Learning rate: 1e-8, 1e-6, 1e-4
- Dropout rate: 0, 0.25, 0.5
- Number of neurons in the initial layer: 32, 64, 128
- Number of neurons in the final layer: 32, 64, 128

In training a classifier, along with the original pair, a flipped version of the pair was also provided. In the flipped version of a pair the upstream window is downstream and the downstream window is upstream. This doubled the number of input pairs provided. LEPAE identified the best-performing combination of hyper-parameters by maximizing the AUROC on the validation pairs.

With the best combination of hyper-parameters for a given window size *r*, pairwise distance *d*, and set of training pairs, LEPAE then trained 5 new classifiers, each with 50,000 training pairs randomly sampled from the training set but with the same combination of hyper-parameters. To compute the LEPAE score for target pairs, which were not in the set of training pairs, LEPAE computed the average probabilities from an ensemble of classifiers. Specifically, for the target pair with pairwise distance *d*, LEPAE applied each of the 5 classifiers trained on pairs with pairwise distance of *d* to obtain the probability of the target pair being classified as a positive pair and also for the flipped pair, resulting in 10 values. In addition, to further reduce variance, for the same target pair of windows with distance of *d*, LEPAE similarly applied 5 classifiers corresponding to a shorter pairwise distance of *d* - *r* to obtain 10 additional values and 5 classifiers corresponding to a longer pairwise distance of *d* + *r* to obtain 10 additional values. The assumption here is that classifiers for adjacent pairwise distances will learn similar information. The final LEPAE score of the target pair was the average of these 30 values, except if the pair had the minimum or maximum pairwise distance, the final score was the average of 20 available values since there is no shorter or longer pairwise distance available, respectively. This procedure of hyper-parameter search, training the final classifiers, and ensemble-based predictions was repeated for each pairwise distance *d*, window size *r*, and set of training pairs.

We used PyTorch (version 0.3.0.post4)^27^ for implementation.

### Defining subset of pairs for training, validation, and test

To generate predictions for all pairs of genomic windows from an odd chromosome or chrX, for a specific pairwise distance *d* and window size *r*, we first randomly selected three even chromosomes (chr6, chr16, chr18) and defined a random subset (*n*=5,000) of pairs of windows from those three chromosomes as a validation set. We selected pairs of windows from the remaining even chromosomes as a training set. To form a test set, we used a random subset (*n*=5,000) of pairs of windows from odd chromosomes.

To generate predictions for all pairs of windows from an even chromosome, we took an analogous approach as above where chr3, chr17, and chr21 were randomly selected for validation. There was no overlap in genomic regions used for training, validation, and test. chrX was excluded from training, validation, and test, but included for prediction and downstream analyses.

### Decision tree evaluation

We trained, applied, and evaluated an ensemble of decision trees using the same procedure as explained above, except we used a decision tree in place of a neural network. We also did hyper-parameter selection as explained above, but for the following set of hyper-parameters unique to decision trees:

- Maximum tree depth: 16, 32, 64, 128, 256
- Minimum fraction of samples required to split at an internal node: 0.0005, 0.001, 0.002, 0.005, 0.01
- Minimum fraction of samples required to be at a leaf node: 0.0005, 0.001, 0.002, 0.005, 0.01

The maximum number of features to consider when looking for the best split was set to square root of the total number of features. As done with neural networks, 10 decision trees were ensembled. We used Scikit-learn (version 0.19.1)^28^ for implementation.

### Filtering pairs

In all analyses, except for reporting predictive performance, all pairs that overlap any assembly gap annotation in either window were excluded since most input features do not map well to these pairs. We downloaded the assembly gap annotation from the UCSC Genome Browser^29^.

### Computing Jaccard index

For each pair, given its two binary input feature vectors, we defined *A* as the set of features set to 1 in the first feature vector and *B* as the set of feature set to 1 in the other feature vector. The Jaccard index between the two vectors was defined as:

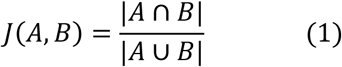

If the numerator was zero, the pair was eliminated from our analysis.

### Hi-C data

We downloaded in situ Hi-C data for GM12878 (experiment 4DNES3JX38V5)^7^ from the 4DN Nucleome Data Portal^24^ from https://4dn-open-data-public.s3.amazonaws.com/fourfront-webprod/wfoutput/a98ca64a-861a-4a8c-92e9-586af457b1fb/4DNFI1UEG1HD.hic. Within the downloaded file, we specifically used values with square root of vanilla coverage (VC_SQRT) normalization applied. We used software straw^30^ to extract the values for the pairs of our interest from the file. If no data was found for a pair in the file, we discarded the pair from our analysis. We downloaded TAD coordinates for GM12878 provided by the 3D Genome Browser^25^ at http://3dgenome.fsm.northwestern.edu/downloads/hg38.TADs.zip.

### Computing mean LEPAE score percentile for pairs of chromatin states

For each state pair, *s_u_* and *s_d_*, its mean LEPAE score percentile was computed as follows. First, to make results comparable with the Hi-C contact frequency and the Jaccard index, the percentile of every score value was computed among pairs of windows with all three scores available. Then, for each pair of windows, *w_u_* and *w_d_*, with at least some portion annotated by states *s_u_* and *s_d_*, respectively, the product of the fraction of bases in window *w_u_* annotated by state *s_u_* and the fraction of bases in window *w_d_* annotated by state *s_d_* was computed. The state pair’s LEPAE score percentile for the window was then multiplied by this product. The overall mean of the state pair was the sum of these weighted score percentiles from all applicable pairs of windows divided by the sum of the products of fractions of bases. We excluded from the analysis pairs of states that together cover less than 0.0005% of pairs of bases. States associated with active transcription, promoters, or enhancers were defined as states belonging to the following state groups, according to how they are annotated in Ref. ^18^: EnhA, PromF, Tx, TxEx, TxEnh, TSS.

### Gene analysis

For each protein-coding gene from GENCODE gene annotation (V38)^23^ with length *l*, we set quartile length *q* to *l* divided by 4. For genes on the positive strand, given the position of the transcription start site (TSS) and transcription end site (TES) of the gene, *s_1_* and *s_2_*, we defined three sets of bases for the gene as follows:

- Upstream of TSS: *s_1_* - 4q, *s_1_* - 3q, *s_1_* - 2q, *s_1_* - q
- Within gene: *s_1_*, *s_1_* + q, *s_1_* + 2q, *s_1_* + 3q, *s_2_*
- Downstream of TES: *s_2_* + q, *s_2_* + 2q, *s_2_* + 3q, *s_2_* + 4q

For genes on the negative strand, the procedure above was reversed such that a distance was subtracted instead of added and vice versa.

We then defined 20 pairs of these bases where at least one base in the pair is within the gene (e.g. *s_1_* - 3q vs. *s_1_* + q) and with a maximum pairwise distance of four quartiles. We then compared two pairs that had the same pairwise distance and shared a base within the gene but one pair crossed a gene boundary, TSS or TES, while the other pair did not (e.g. *s_1_* - q vs. *s_1_* + q compared to *s_1_* + q vs. *s_1_* + 3q). The LEPAE score for a pair of bases was the score of the two 1-kb windows that overlap the bases. This excluded pairs of bases less than 1 kb apart from the analysis.

This procedure allowed us to evaluate whether the LEPAE score, Jaccard index, or Hi-C contact frequency favors pairs of windows within genes over those crossing gene boundaries. Using quartile lengths of each gene rather than a fixed pairwise distance allowed us to control for varying gene lengths.

### Fine-mapping analysis

We obtained fine-mapped variants for 94 UK Biobank traits from https://www.finucanelab.org/data^21,22^. For a pair of variants in windows *i*, *j* where we assume *i* < *j* and they are *d* bases apart, we added as controls pairs of positions corresponding to (*i - d, i*) and (*j*, *j + d*). We conducted the analysis at 1-kb resolution considering variants within 100 kb and also at 10-kb resolution considering pairs of variants within 1 Mb. For each resolution, we separately analyzed the fine-mappings based on FINEMAP^20^ and SuSiE^19^. For each resolution and fine-mapping method, we computed the average LEPAE score for the original pairs of variants and control pairs for each of the 94 traits and determined which average was greater. We repeated the analysis for the GM12878 Hi-C contact frequency and Jaccard Index.

## Author contributions statement

S.K. and J.E. developed the method, analyzed the results, and wrote the paper.

## Acknowledgments

We thank the ENCODE and Roadmap Epigenomics consortia for generating data and making it publicly available. We thank members of the Ernst lab for useful discussions. We acknowledge funding from US National Institutes of Health (DP1DA044371, U01MH130995, and U01HG012079 to J.E.), a Rose Hills Innovator Award (J.E.), and a UCLA Dissertation Year Fellowship (S.K.).

