## Supplementary Figures 1-7 for "Learning a Pairwise Epigenomic and Transcription Factor Binding Association Score Across the Human Genome"

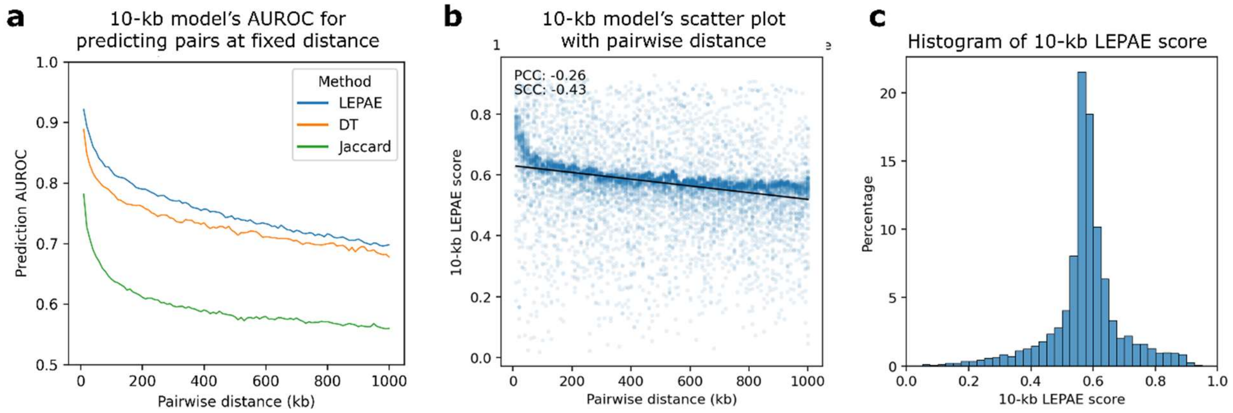


**Supplementary Figure 1. Characteristics of the 10-kb LEPAE score.**

**a.** Relationship between pairwise distance and prediction AUROC for 10-kb model. For each pairwise distance (x-axis), mean prediction AUROC of the 10-kb LEPAE score for distinguishing pairs of windows at that distance from randomly mismatched pairs of the same windows is shown in blue. The mean is computed from two sets of classifiers trained on non-overlapping training sets (**Methods**). Mean AUROC values are shown when a decision tree (DT) instead of a neural network was used as the supervised classifier are shown in orange (**Methods**). Mean AUROC values computed using Jaccard index instead of the LEPAE score to perform the same classification task are shown in green. Values belonging to the same method are connected by piecewise linear interpolation.

**b.** Scatter plot showing with a blue dot for each pair of windows the pairwise distance (x-axis) and the 10-kb LEPAE score (y-axis). Ten thousand random pairs are shown. A linear regression line fitted to the ten thousand random pairs is shown in black. PCC and SCC, computed from 1 million randomly selected pairs, are shown in the top left.

**c.** Distribution of the 10-kb LEPAE score. Forty bins ranging from 0 to 1 with increments of 0.025 were used.

A 1-kb version of this figure is in **Figure 3**.

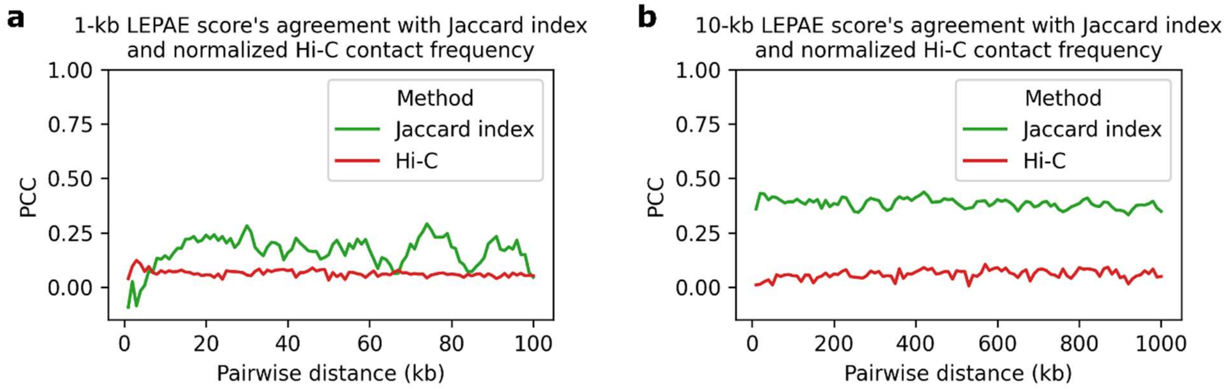


**Supplementary Figure 2. LEPAE score's relationship to Hi-C contact frequency and Jaccard index between input features**

**a.** Shown for each pairwise distance (*x*-axis) is PCC of the LEPAE score with either Jaccard index (green) or normalized Hi-C contact frequency (red) for pairs of windows with the specified distance between them. Values belonging to the same method are connected by piecewise linear interpolation.

**b.** Same as **a** but for 10-kb LEPAE score instead of 1-kb LEPAE score.

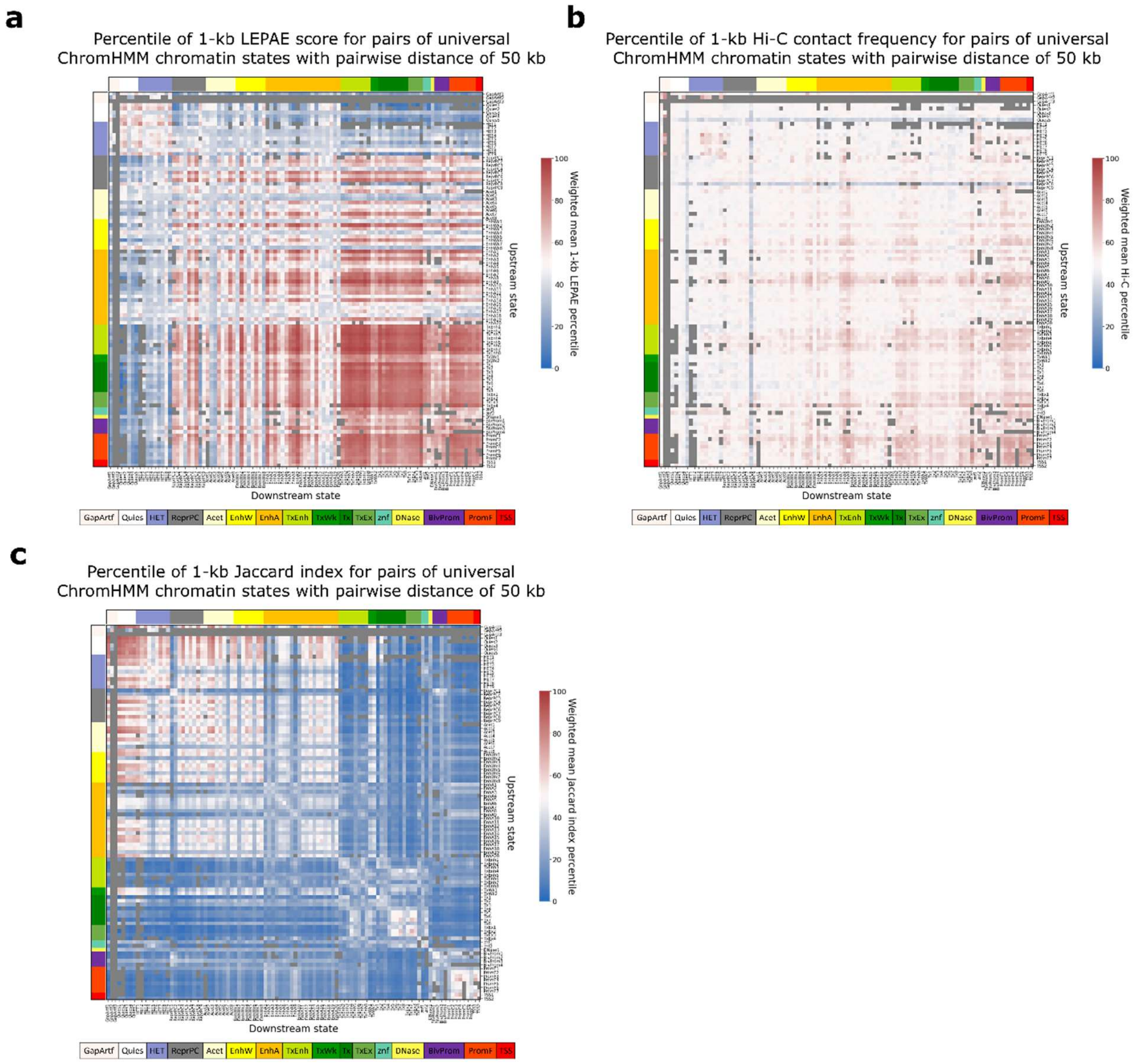


**Supplementary Figure 3. Heatmaps of mean percentile of 1-kb LEPAE score, Hi-C contact frequency, and Jaccard index for pairs of chromatin states with pairwise distance of 50 kb**

**a.** Each cell in the heatmap corresponds to a state pair, one annotating the upstream window of a pair of 1-kb windows (row) and the other annotating the downstream 1-kb window of the same pair (column) with their pairwise distance fixed to 50 kb. The states are from a universal chromatin state annotation based on more than 1000 epigenomic datasets from more than 100 cell or tissue types (Vu and Ernst, 2022). The ordering of states in the rows and columns are the same. Color shown next to the topmost row or leftmost column corresponds to the state group of each state along the column or row, respectively, according to the legend on the bottom left. The colors shown in the cells correspond to a weighted mean LEPAE score percentile of pairs of windows that are 50 kb apart and are annotated by the states specified in the row and column (**Methods**). Color legend for the score is shown on the right. A similar version of this figure but for pairwise distance of 5 kb is in **Figure 5a**.

**b.** Similar to **a** but with each cell showing percentile based on Hi-C contact frequency instead of LEPAE score

**c.** Similar to **a** but with each cell showing percentile based on Jaccard index instead of LEPAE score

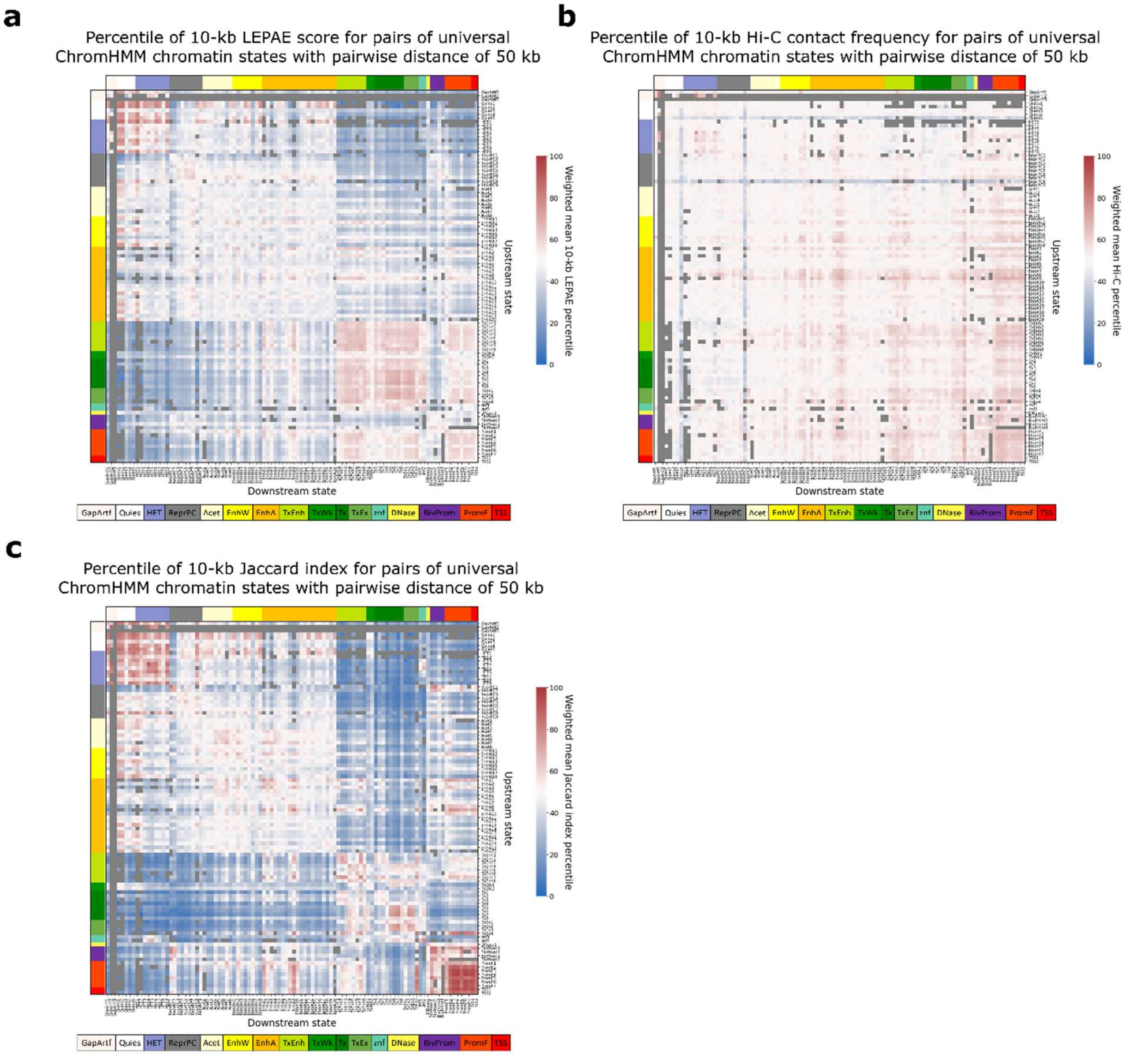


**Supplementary Figure 4. Heatmaps of mean percentile of 10-kb LEPAE score, Hi-C contact frequency, and Jaccard index for pairs of chromatin states with pairwise distance of 50 kb**

Similar to **Supplementary Fig. 3** but showing results from analyzing at a 10-kb score resolution and with pairwise distance of 50 kb

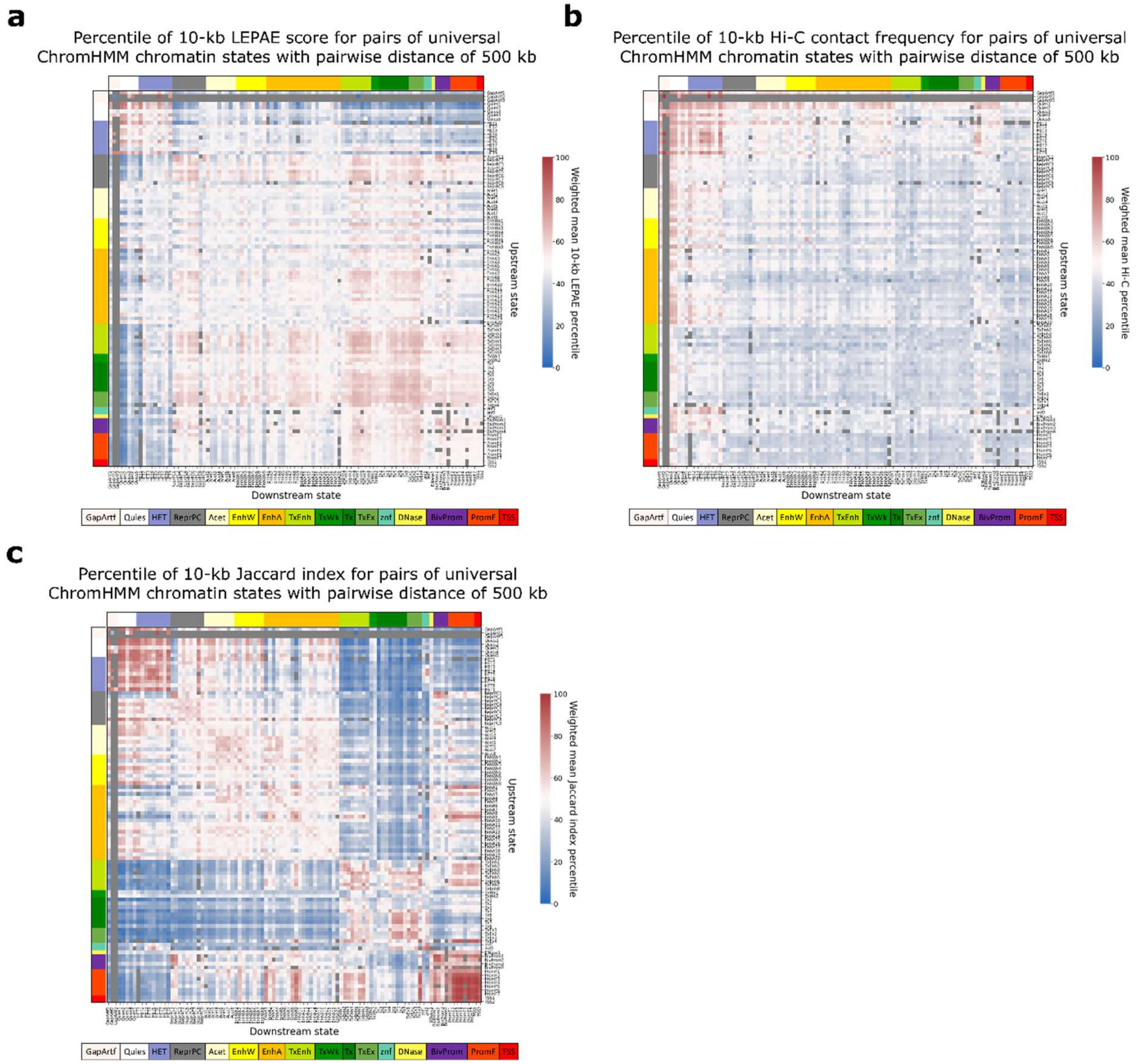


**Supplementary Figure 5. Heatmaps of mean percentile of 10-kb LEPAE score, Hi-C contact frequency, and Jaccard index for pairs of chromatin states with pairwise distance of 500 kb**

Similar to **Supplementary Fig. 3** but showing results from analyzing at a 10-kb score resolution and with pairwise distance of 500 kb

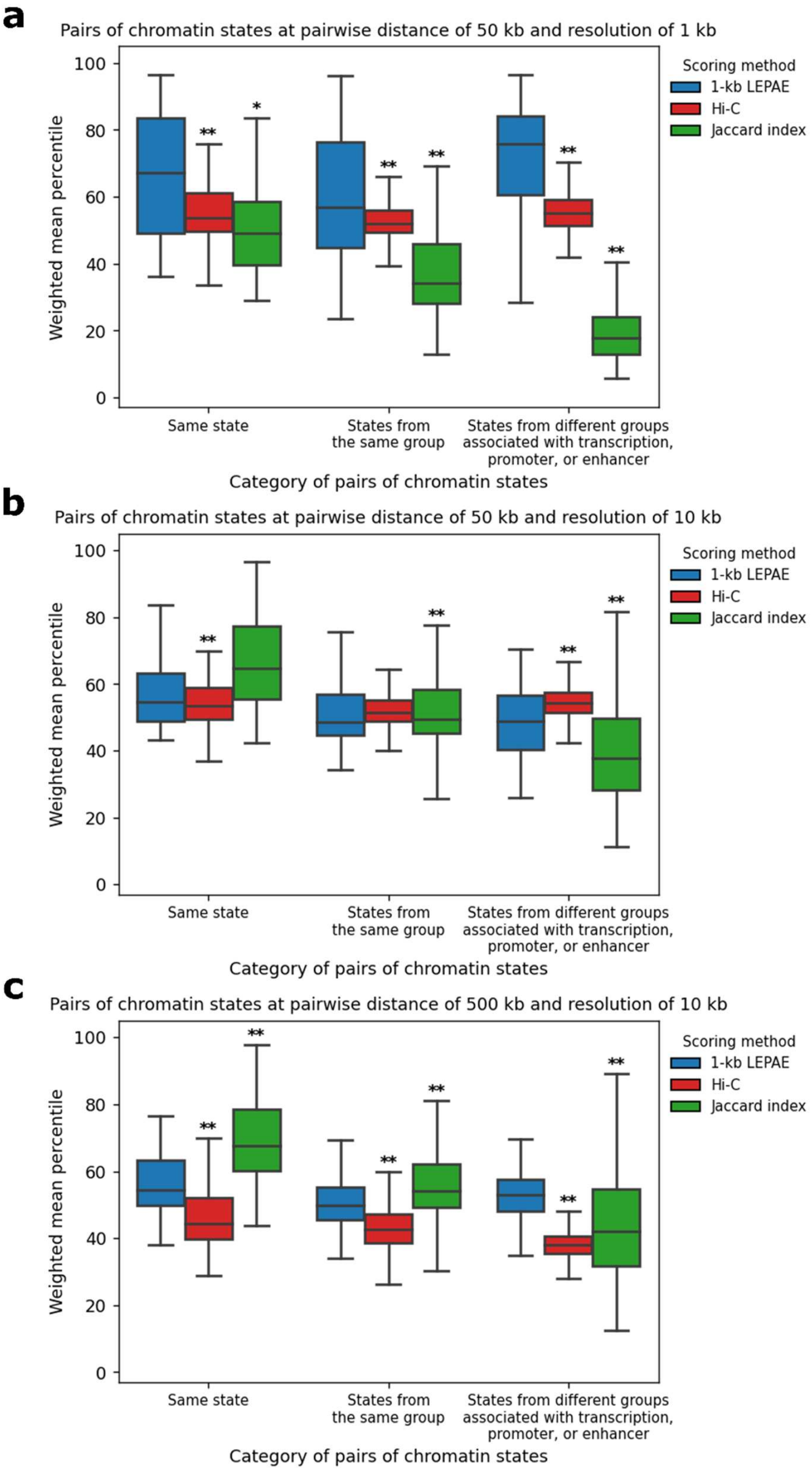


**Supplementary Figure 6. Distribution of weighted mean percentile of different scores for pairs of chromatin states.**

**a**. Shown for three different categories of pairs of ChromHMM chromatin states are the distribution of the weighted mean percentile of the 1-kb LEPAE score, Hi-C contact frequency, or Jaccard index, at a fixed pairwise distance of 50 kb as done in **Supplementary Fig. 3**. Among the three categories, the first category shown on the leftmost position on the x-axis corresponds to pairs of the same state. The second, shown in the middle, corresponds to pairs of different states belonging to the same state group. The last category shown on the rightmost position corresponds to pairs of states where both are a transcription, promoter, or enhancer associated states but belong to different state groups. These three categories of pairs of states do not overlap with each other. Within each category, the distributions of the weighted mean percentile of the 1-kb LEPAE score, Hi-C contact frequency, and Jaccard index are shown in blue, red, and green, respectively, according to the score legend on the right. One asterisk and two asterisks above a distribution denotes that there is a significant difference between it and the distribution of weighted 1-kb LEPAE score percentiles within the same category based on a Mann-Whitney U test for p-values less than 0.001 and 0.0001, respectively. Similar versions of this figure but for pairwise distance of 5 kb is shown in **Figure 5d**.

**b.** Similar to **a** but showing results from 10-kb score resolution at a pairwise distance of 50 kb, as done in **Supplementary Fig. 4.**

**c.** Similar to **a** but showing results from 10-kb score resolution at a pairwise distance of 500 kb, as done in **Supplementary Fig 5.**

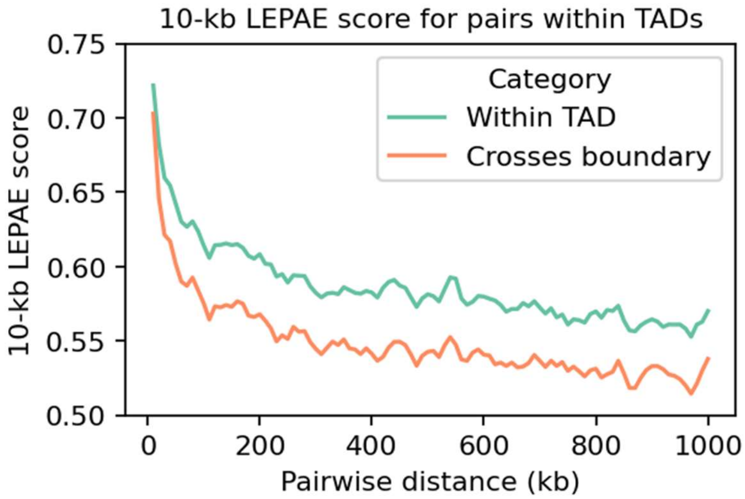


**Supplementary Figure 7. 10-kb LEPAE score's relationship to TAD annotation**

Shown for each pairwise distance (x-axis) is the mean 10-kb LEPAE score for pairs of windows located within a topologically associating domain (TAD) (turquoise) or the mean score for pairs of windows crossing a TAD boundary (peach). Values belonging to the same category are connected by piecewise linear interpolation. A 1-kb score resolution version of this figure is in **Figure 6**.
